# Impact of APOE ε3 and ε4 genotypes on plasma proteome signatures in Alzheimer’s disease

**DOI:** 10.1101/2022.01.29.478291

**Authors:** Gurjeet Kaur, Anne Poljak, Colin L Masters, Christopher Fowler, Perminder Sachdev

**Affiliations:** Centre for Healthy Brain Ageing, School of Psychiatry, University of New South Wales, Sydney, NSW, 2052, Australia; Mark Wainwright Analytical Centre, Bioanalytical Mass Spectrometry Facility, University of New South Wales, Sydney, NSW, 2052, Australia; Florey Institute, The University of Melbourne, Victoria, Australia; Neuropsychiatric Institute, Euroa Centre, Prince of Wales Hospital, Sydney, NSW, 2052, Australia

**Keywords:** Plasma, proteomics, *APOEε3*, *APOEε4*, Alzheimer’s disease

## Abstract

The ε*4* allele of the apolipoprotein E (*APOE*) gene is a high-risk factor for Alzheimer’s disease (AD). However, approximately 25%–40% of patients with AD do not carry the *APOEε4* allele, and the pathophysiological mechanisms underlying AD are less evident in these individuals. The main objective of this study was to understand better the changes in plasma that may contribute to disease pathogenesis in AD and how *APOEε3* and *APOEε4* contribute to biomarker profiles in AD. We conducted an in-depth plasma proteomics analysis using intensive depletion of high-abundant plasma proteins using the Agilent multiple affinity removal liquid chromatography (LC) column-Human 14 (Hu14) followed by sodium dodecyl sulphate-polyacrylamide gel electrophoresis (SDS PAGE) technique. In this study, we identified a high number of protein expression alterations in plasma which were found uniquely in *APOEε3* and *APOEε4* carriers. These differentially expressed proteins (DEPs) were associated with several molecular functions, including complement cascade, glycolysis, metabolism, plasma lipoprotein assembly, remodelling, and clearance. In addition to unique changes in both *APOE* genotypes, many proteins were also dysregulated in the presence of both *APOEε3* and *APOEε4* genotypes depicting the involvement of these proteins in the pathogenesis of AD regardless of the *APOE* genotypes. We also compared the plasma proteomes of *ε4* and *ε3* carriers in normal controls, which provided insight into factors that may provide protection from progression to AD despite the presence of the *ε*4 allele. Furthermore, our findings also identified some proteins previously discovered in AD CSF and brain proteomics signatures that could provide clinically meaningful information.

## Introduction

One of the barriers to developing effective therapies for Alzheimer’s disease (AD), the most common cause of dementia, lies in the lack of a comprehensive understanding of the brain mechanisms leading to neurodegeneration. One key knowledge gap is in understanding how genetic risk factors contribute to disease pathogenesis. There are numerous genetic risk factors for developing sporadic AD, the strongest of which is the apolipoprotein E epsilon 4 allele inheritance (*APOEε4*)^1-4^. Three alleles, i.e., *APOEε2, APOEε3*, and *APOEε4*, result in six possible genotypes (*APOE* 2/2, 2/3, 3/3, 2/4, 3/4, and 4/4). These three polymorphic alleles, i.e., ε2, ε3, and ε4, have a worldwide frequency of 8.4%, 77.9%, and 13.7%, respectively^1,5^. Recent studies reported that approximately 65% of individuals with late-onset familial and sporadic AD bear the *APOEε4* allele^6^. One copy of *APOEε4* is associated with a threefold increase in disease risk, while two copies are associated with a more than tenfold increase in risk^7^.

Emerging data suggest that *APOEε4 is* involved in several functions, including metabolism, neuroinflammation, impaired amyloid clearance, transport, synaptogenesis, and glucose, lipid, and cholesterol metabolism in the brain^8,9 10^. In animal and cellular models, *APOEε4* has been linked to decreased cellular plasticity^11^. In addition, *APOE* plays a critical role in lipid transport and cholesterol homeostasis in the brain, as it does peripherally ^1,12^. In the central nervous system (CNS), *APOE* is mostly expressed in astrocytes, and it facilitates the transportation of cholesterol to neurons by binding to LDLR family members, known as APOE receptors. *APOEε4* has been found to be hypolipidated and less effective at inducing cholesterol efflux than *APOEε3*, implying that the pathological effects of *APOE* may be associated with lipid metabolism^3^. However, approximately 25%–40% of patients with AD do not carry the *APOEε4* allele, and the pathophysiological mechanisms underlying AD are less clear in these individuals^1,13^.

Unbiased proteomics analysis permits the simultaneous evaluation of many molecular processes in patients. To explore this, several research studies have used a CSF proteomics technique and described protein signatures linked with AD across the cognitive range^14,15^. Proteomics investigations on readily available fluids such as serum/plasma, on the other hand, are underutilized. To gain a better understanding of how *APOE* genotypes may influence AD pathology, we used a plasma proteomic approach to test the hypothesis that protein signatures can be detected that show *APOE* genotype-dependent associations with AD. The main objective of this study was to better understand the changes in plasma that may contribute to disease pathogenesis in AD and how *APOEε3* and *APOEε4* contribute to biomarker profiles in AD. In this study, AD and cognitively normal age-matched control carriers of the *APOEε4* allele and AD and control homozygous *APOEε3* carriers were included. Furthermore, all AD (whether *ε*3 or *ε*4) were Pittsburgh compound B (PiB) positron emission tomography (PET) positive (high or very high), whereas all controls (whether *ε*3 or *ε*4) were PiB PET negative.

## Materials and Methods

### Cohort and samples

Plasma samples were obtained from the Australian Imaging, Biomarker & Lifestyle Flagship Study of Ageing (AIBL) from participants aged 70-90 years^16^. The University of Melbourne Human Research Ethics Committee approved the collection of the AIBL cohort, while the UNSW Human Research Ethics Committee approved the current study. All work complied with the Declaration of Helsinki guidelines.

In total, we profiled 40 human plasma samples using label-free proteomics in the following four groups: **1**. *APOE* ε4/ε3 carriers without AD are denoted as <u>CNTLE4</u>, **2**. *APOE* ε4/ε4 carriers with AD symptoms denoted as <u>ADE4</u> **3**. *APOE* ε3/ε3 carriers without AD denoted as <u>CNTLE3</u> and **4**. *APOE* ε3/ε3 carriers with AD symptoms are denoted as <u>ADE3</u>.

### Depletion of high abundant proteins using Human 14 (Hu14) immunoaffinity Columns

The protocol for removing plasma high abundance proteins followed by fractionating the low abundance proteins was adapted from a previously published approach^17^. The top 14 high-abundance plasma proteins (albumin, IgG, antitrypsin, IgA, transferrin, haptoglobin, fibrinogen, α-2-macroglobulin, α-1-acid glycoprotein, IgM, apolipoprotein AI, apolipoprotein AII, complement C3, and transthyretin) were depleted using an Hu14 column (4.6 × 100 mm, Agilent). The plasma (50 µL) was diluted with 150 µL of buffer A (1:4 dilutions, as recommended by Agilent Technologies), and then filtered to remove particulates using a 0.45 μm spin filter (Spin-X centrifuge tube filter, 0.45 μm Cellulose Acetate, Merck, Germany). Samples were then injected (100 µL) onto the Hu14 column. Chromatography and fraction collection was performed on an Agilent 1290 UPLC system (Agilent, Santa Clara, CA) with Hu14 buffers A and B purchased from Agilent (Santa Clara, CA), and manufacturer’s instructions followed for protein binding and elution (Agilent, Santa Clara, CA). The low abundance protein fraction was further fractionated by 1D SDS PAGE and analyzed using LC-MS/MS. Participant demographics are shown in Table 1. Each sample consisted of ten SDS PAGE fractions, so a total of 400 LC-MSMS runs were performed to ensure adequate coverage of the plasma proteome.

**Table 1:** Participant demographic details used for this study.

| **Total participants** | **Control**<br>***APOE*ε3** | **Control**<br>***APOE*ε4** | **AD**<br>***APOE*ε3** | **AD**<br>***APOE*ε4** | **Kruskal-Wallis**<br>**statistic** | **Kruskal-Wallis P**<br>**value** |
| --- | --- | --- | --- | --- | --- | --- |
| Total participants in each wave | 10 | 10 | 10 | 10 | NA | NA |
| Age in years mean±SD<br>(CV%) | 72.40±5.0<br>(6.9%) | 72.10±6.2<br>(8.6%) | 70.50±5.6<br>(8.03%) | 70.70±6.5<br>(9.28%) | 0.42 | 0.93 |
| Education (years) | 12.90±3.2<br>(24.91%) | 14.80±2.3<br>(15.86%) | 11.60±3.1<br>(27.02%) | 12.70±2.2<br>(17.43%) | 7.47 | 0.05 |
| Sex (n) | F=5, M=5 | F=6, M=4 | F=6, M=4 | F=5, M=5 | NA | NA |
| *APOE* status | E3/3 | E3/4 | E3/3 | E3/4 | NA | NA |
| MMSE<br>mean±SD (CV%) | 29.00±1.0<br>(3.63%) | 29.50±0.5<br>(1.78%) | 19.50±4.9<br>(25.5%) | 21.40±7.6<br>(35.8%) | 30.51 | 0.00 |
| Hypertension in number of<br>participants (%) | 50% | 40% | 60% | 30% | NA | NA |
| Diabetes in number of participants<br>(%) | 10% | 10% | 0% | 10% | NA | NA |
| Cholesterol (mmol/L)<br>mean±SD (CV%) | 5.29±1.2<br>(22.81%) | 5.45±1.5<br>(27.83%) | 5.65±1.0<br>(18.34%) | 5.63±1.1<br>(20.83%) | 0.53 | 0.91 |
| Triglyceride (mmol/L)<br>mean±SD (CV%) | 1.02±0.2<br>(26.06%) | 1.43±0.52<br>(36.72%) | 1.23±0.50<br>(40.50%) | 1.63±1.45<br>(89.25%) | 3.96 | 0.26 |
| HDL-Chol (mmol/L)<br>mean±SD (CV%) | 1.72±0.53<br>(31.03%) | 1.52±0.48<br>(32.07%) | 1.67±0.42<br>(25.10%) | 1.47±0.37<br>(25.69%) | 0.61 | 1.81 |
| LDL-Chol (mmol/L)<br>mean±SD (CV%) | 3.09±0.93<br>(30.38%) | 3.27±1.21<br>(37.06%) | 3.41±0.94<br>(27.64%) | 3.40±1.20<br>(35.48%) | 0.65 | 0.88 |
| Urea (mmol/L)<br>mean±SD(CV%) | 6.40±1.17<br>(18.43%) | 5.98±8.90<br>(21.89%) | 5.36±0.92<br>(17.18%) | 6.27±2.09<br>(33.14%) | 3.69 | 0.29 |

SD= standard deviation; cv= coefficient of variations

### Fractionation of low abundance proteins using 1D-SDS PAGE

We followed a previously published procedure for SDS-PAGE, band cutting, trypsin digestion, sample preparation, and mass spectrometric analysis^17^. Equal amounts of total protein (50 µg) from the Hu14 depleted plasma were filtered using Amicon ultra 3kDa centrifugal filter units (MERCK, New Jersey, USA), dried in speed vac (ThermoFisher, Massachusetts, USA), and reconstituted to a final volume of 20 µL by adding 5 µL LDS sample buffer Invitrogen NuPAGE (ThermoFisher, Massachusetts, USA), 2 µL reducing agent Invitrogen NuPAGE (ThermoFisher, Massachusetts, USA), and 13 µL deionized water (MilliQ). After briefly heating samples (10 minutes, 70°C), they were separated by 1D SDS/PAGE using Invitrogen NuPAGE 4-12% Bis-Tris midi gels (ThermoFisher Scientific, Massachusetts, USA) and Invitrogen MES running buffer using the manufacturer’s instructions (ThermoFisher Scientific, MA, USA). The gel was then stained using colloidal coomassie G250 (Figure S1). The protein lanes were cut into gel bits the following destaining using a 24-band lane cutting blade. The gel bands were concatenated into ten vials for in-gel trypsin digestion, peptide recovery, and label-free LCMSMS quantification. A total of 10 biological replicates (subjects) were used per group, i.e., ADE4, CTRLE4, ADE3, CTRLE3.

### Computational Analysis

Two search engines were used to analyze the raw files, including ProteomeDiscoverer v2.4 (Thermo Fisher Scientific, Waltham, MA) and Scaffold Q+ software v 4.11.0 (Proteome Software, Portland, OR). A minimum of two unique peptides per protein were prerequisites for protein identification and quantitation for both data analysis software. In conjunction with the reversed decoy and frequent contaminant sequences, all search engines used the UniProt Homo sapiens (human) database for MS and MS/MS spectral mapping. Five parts per million (5ppm) mass tolerance was used to match peaks to theoretical ion series. The false discovery rate (FDR) was set to 1% to ensure that only highly confident protein identifications were made. Trypsin was chosen as the specific enzyme, with a maximum of two missed cleavages. Variable acetylation at the N-terminus of proteins, methionine oxidation, and fixed carbamidomethylation of cysteines was used for all database searches. All parameters were kept consistent between the two search engines. The software used enabled the simultaneous application of two distinct approaches for label-free quantification: peak area integration (PD2.4) and spectral counting (Scaffold).

Each AD group required independent proteomics data processing using the PD2.4 and Scaffold search engines. In supplementary Figure S2, scatter plots and regression analysis comparing PD2.4 versus scaffold fold-change are displayed. To identify proteins with significant expression differences between groups, we used the following inclusion criteria: proteins quantified in >5 individuals, proteins identified with a minimum of two peptides per protein, and consistent direction of protein fold change across two bioinformatics platforms using orthogonal quantification approaches (peak area ratio with PD2.4 and spectral counting with Scaffold) with a fold change of at least 20% (≤0.08 and ≥ 1.2) in both search engines are summarized in Table S2 for ADE4/CTRLE4, Table S3 ADE3/CTRLE3, Table S4 CTRLE4/CTRLE3 and Table S5 for ADE4/CTRLE3.

### Bioinformatics Analysis

Bioinformatics analyses were performed using RStudio version 1.2.5033 and R version 3.6.3 to create heatmaps and volcano plots, using the heatmap function and ggplot2 package. Gene ontology and enrichment plot analysis were performed using Bioconductor’s GOstats and DOSE package. Results from the gene ontology analysis were only studied if more than two genes from the experimental data set were included with a particular term. Volcano plot analysis was performed using the Enhanced Volcano package from Bioconductor^18^. Venn diagrams were plotted using Venny 2.114. We used differentially expressed proteins (DEPs) to compare biological processes and pathways affected in AD versus control using gene ontology (GO) enrichment analysis on the Database for Annotation, Visualization, and Integrated Discovery (DAVID) v6.8. Additionally, STRING (11.0) explored gene interaction and co-expression patterns for differentially expressed genes (DEGs).

## Results

### Sample characteristics by *APOEε3* and *APOEε4* genotype

We profiled 40 human plasma samples using label-free proteomics in the following four groups: **(1)** *APOE* ε4/ε3 carriers, cognitively normal controls with negative PiB PET denoted as CNTLE4 **(2)** *APOE* ε4/ε4 carriers with AD symptoms and positive PiB PET denoted as ADE4 **(3)** *APOE* ε3/ε3 carriers, cognitively normal controls with negative PiB PET denoted as CNTLE3 **(4)** *APOE* ε3/ε3 carriers with AD symptoms and positive PiB PET denoted as ADE3. We identified 1,055 proteins (false discovery rate <1%) with 23,242 total peptides using Proteome Discoverer 2.4 (PD2.4) search engine and 800 proteins using Scaffold (Table S1). More than 700 identified proteins were common in both software techniques.

An overview of the study populations and proteomic workflow is shown in Figure 1A. Box plots show the similar distribution and protein abundance variation across all 40 plasma samples (Figure 1B). The overall similarity of low abundance proteins across samples is also evident by SDS PAGE (Supplementary Figure S1). Unsupervised hierarchical clustering analysis (HCA) of grouped abundances (data from PD2.4 software processing) is presented in Figure 1C. It shows that control and AD samples of both ADE3 and ADE4 carriers cluster together more closely based on diagnosis sample type (i.e., control vs. AD) rather than *APOE* allele type. Nevertheless, distinct proteomic profiles are observed in each of the four groups since all heat maps are quite distinct (Figure 1C). The scatter plot depicting AD and CTRL data points analyzed on both PD2.4 and Scaffold is shown in Figure 1D. The detailed scatter plots and density plots were plotted using the complete list of proteins from both the search engines in Figure S2i. In Figure S2ii, the scatter plots show only the differentially expressed proteins (DEPs) using all the analyses, i.e., **A**. ADE3/CTRLE3, **B**. ADE4/CTRLE3, **C**. ADE4/ADE3, **D**. CTRLE4/CTRLE3, and **E**. ADE4/CTRLE4. The DEPs with a similar direction of change using both orthogonal quantification techniques, PD2.4 peak area ratio, and scaffold spectral counting are shown in Figure 1E. The bar graph shows the total number of proteins upregulated and downregulated in five comparisons, i.e., ADE3/CTRLE3, ADE4/CTRLE3, CTRLE4/CTRLE, ADE4/CTRLE4, and ADE4/ADE3 (Figure 1E).

**Figure 1:**
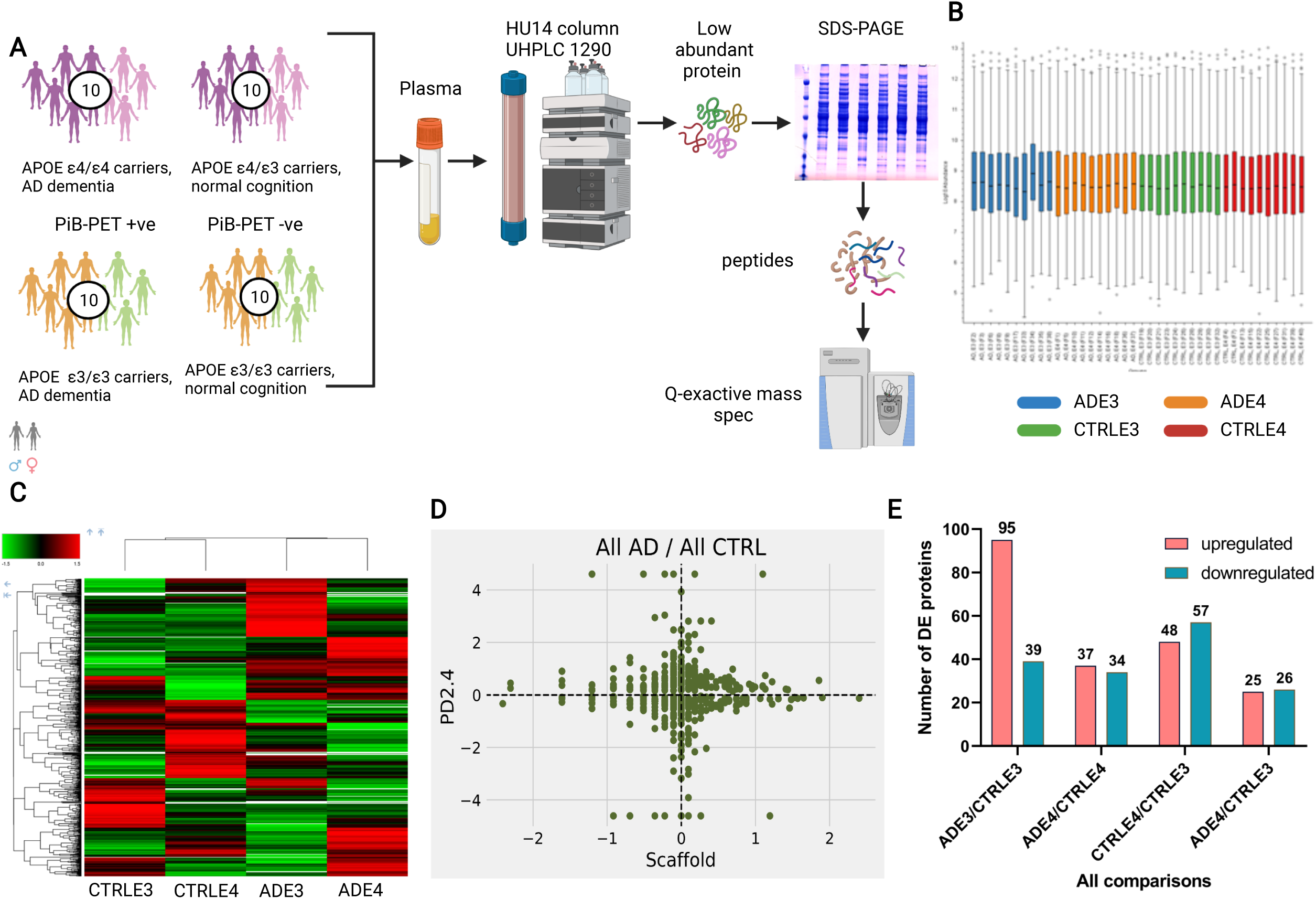
Workflow of plasma proteome profiling and comparison of APOEε3 and APOEε4 genotype. Overview of the study populations and schematic of the proteomic workflow. Dark and light shades represent male and female subjects, respectively. The flow diagram outlines steps of sample preparation through to data acquisition. Box-and-whisker plots of abundance values of all 40 individual samples. The small horizontal line within each box denotes the median value, and the upper and lower ranges (whiskers) indicate the 5 and 95 percentiles of the abundance values, respectively (output from ProteomeDiscoverer 2.4 software). Hierarchical clustering analysis (HCA) of the grouped proteomes of 40 individuals; 10 individuals in each category, i.e., ADE3, control ε3, ADE4, and control ε4 (output from ProteomeDiscoverer 2.4 software). Scatter plot of all protein abundance data (control and AD) comparing PD2.4 and Scaffold. Scatter plots stratified by specific data subsets are shown in Supplementary Figure S2. Global analyses of proteomic changes in specific AD and control comparison groups. Bar graph showing the total number of proteins upregulated (pink) and downregulated (blue) in ADE3/CTRLE3, ADE4/CTRLE4, CTRLE4/CTRLE3, ADE4/CTRLE3, ADE4/ADE3. The numbers at the top of each bar indicate the number of differentially expressed proteins (DEPs) in that category. This data was based on the criteria for DEP selection outlined in the method (i.e., only high confidence protein identifications are used, each identified with a minimum of 2 unique peptides, ≥20% fold-change in group comparisons, the consistent direction of fold change using two orthogonal quantification methods, change identified in >5 individuals per group).

### Overall plasma proteome changes in AD vs. controls in *APOEε3* carriers

A heatmap of the total of 134 proteins that were differentially expressed (95 upregulated and 39 downregulated) in AD *ε3* relative to *ε3* controls (ADE3/CTRLE3) is shown in two panels (Figure 2Ai and 2Aii) for better visibility of the protein acronyms and fold changes. The DEPs with the highest fold change in the ADE3/CTRLE3 group are shown in a volcano plot, using the PD2.4 abundance ratios (Figure 2B), with the complete list of DEPs shown in Table S3 and Table 2. A subset of 65 DEPs (48 upregulated and 17 downregulated) was unique to the ADE3/CTRLE3 group and did not exhibit differential expression in other comparison groups, including; ADE4/ADE3, ADE4/CTRLE4, or CTRLE4/CTRLE3 comparisons (Figure 3B and Table 2).

**Table 2:**
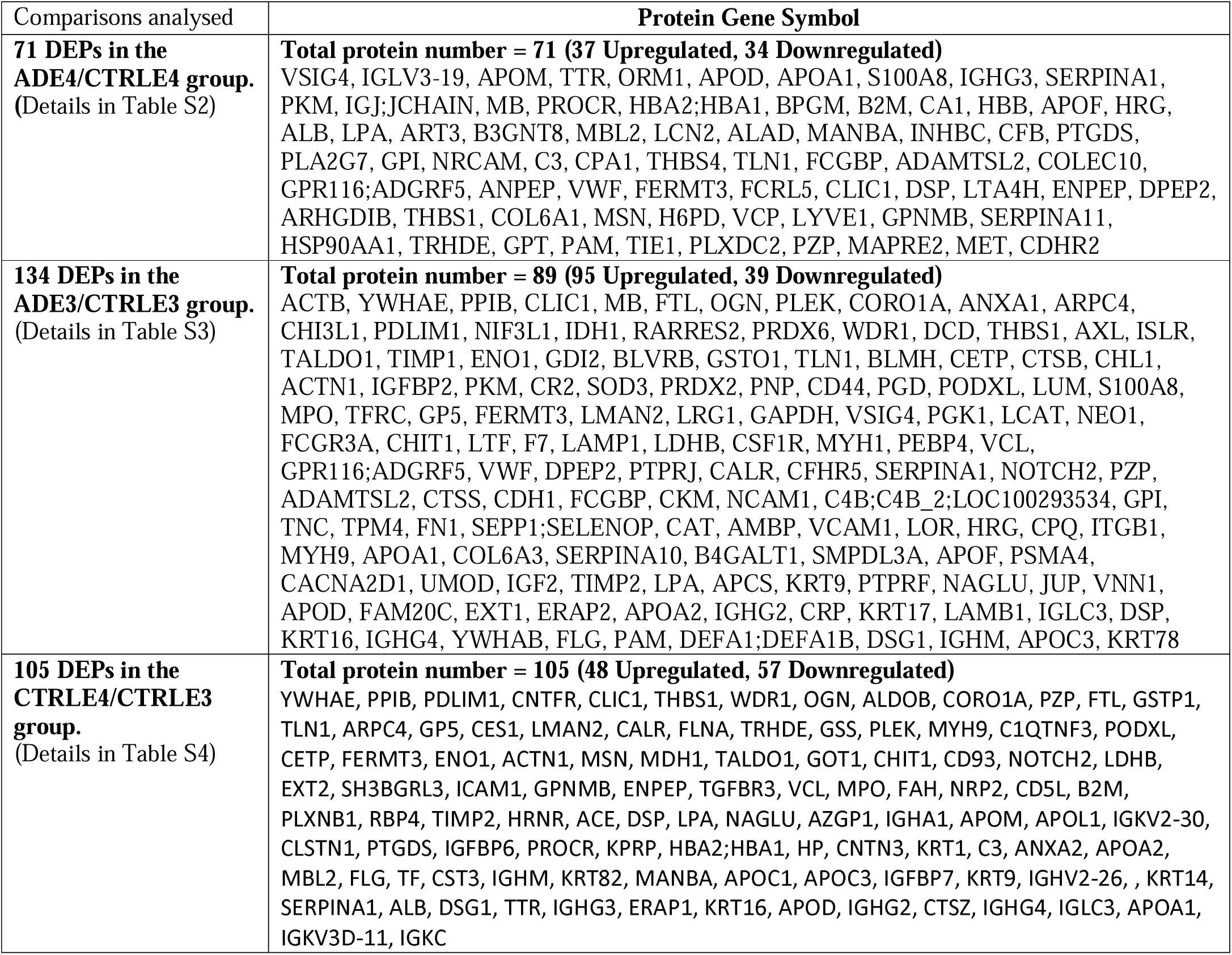
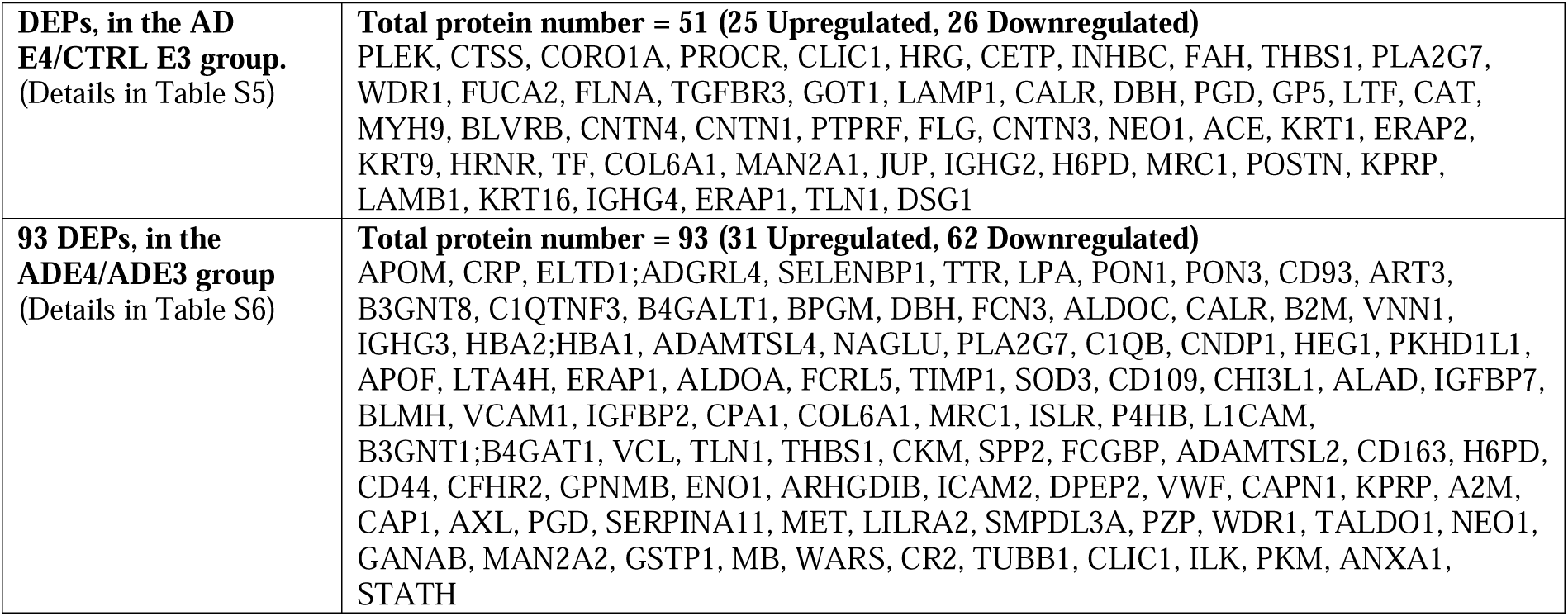
The final list of differentially expressed proteins (DEPs) in all the comparisons analysed. This list contains DEPs quantified in >5 individuals, proteins identified with a minimum of two peptides/protein, consistent direction of protein fold change across two bioinformatics platforms with orthogonal quantification approaches (peak area ratio with PD2.4 and spectral counting with Scaffold) with a fold change of at least 20% (≤0.08 and ≥ 1.2) in both search engines.

**Figure 2:**
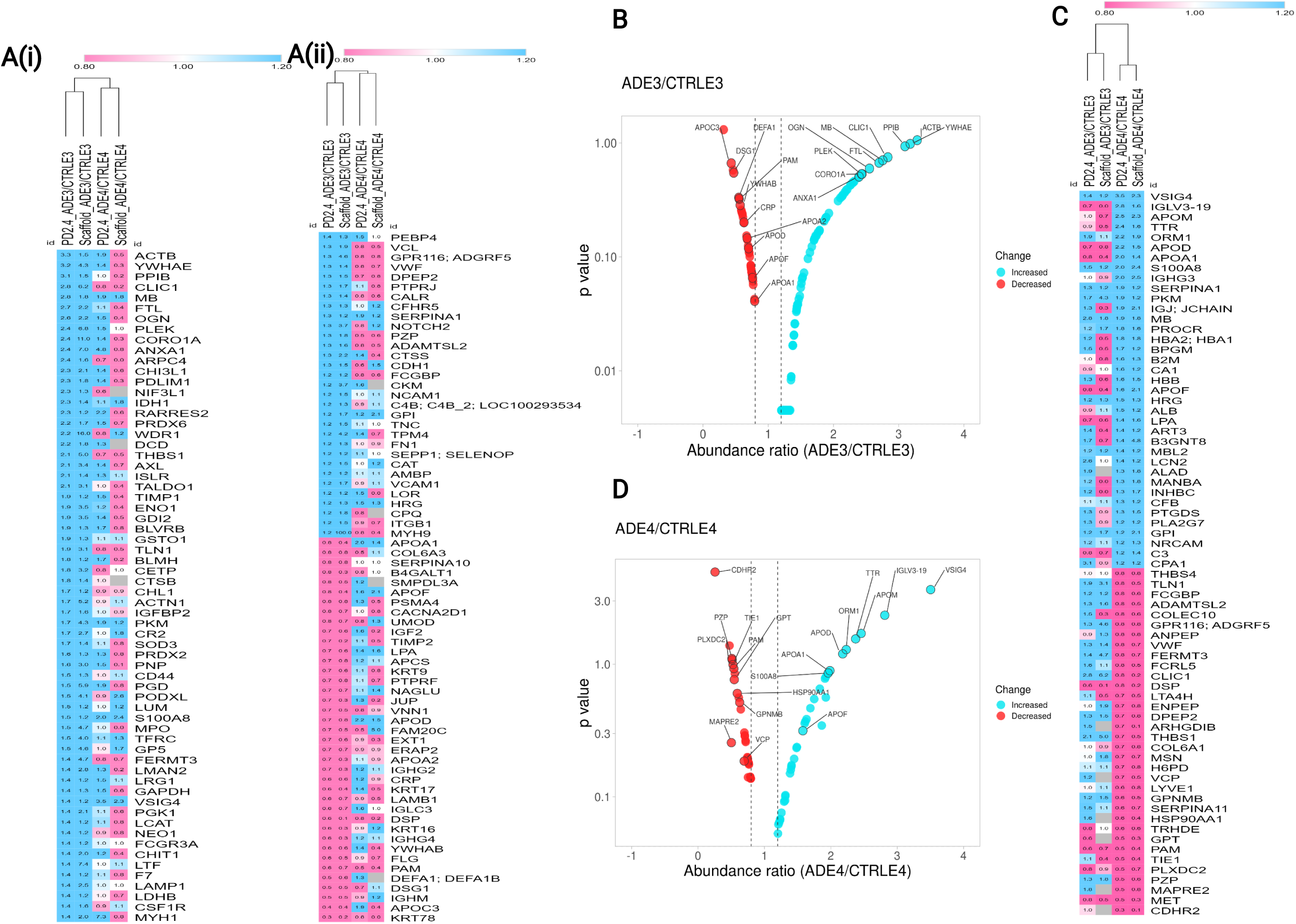
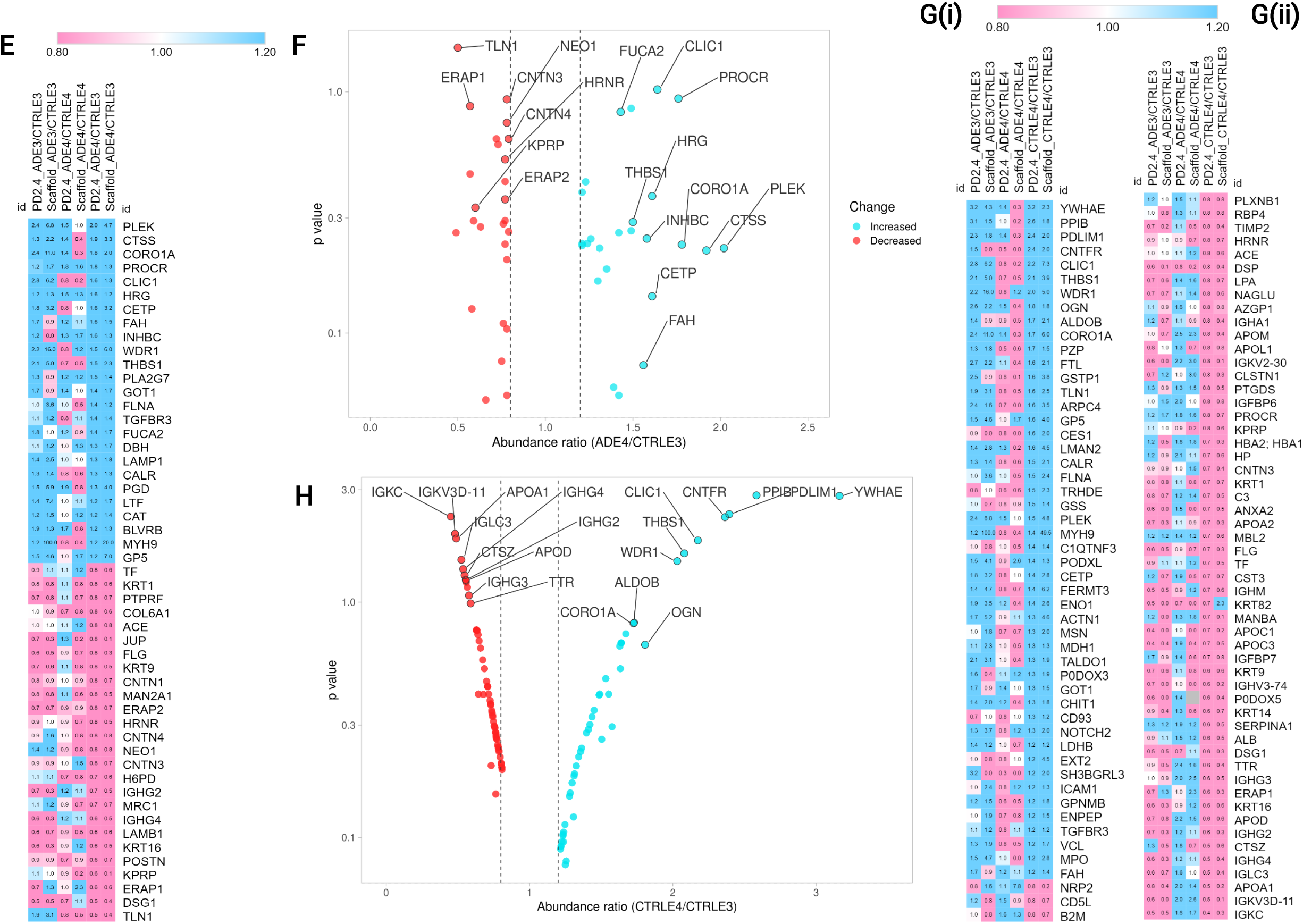
Global analyses of proteomic changes in ADE3/CTRLE3 and ADE4/CTRLE4 analysis. **Ai and ii**. Heatmap showing a total of 134 proteins differentially expressed, including 95 upregulated and 39 downregulated DEPs in ADE3 relative to E3 controls based on the ADE3/CTRLE3 data shown in Table S3. The heatmap is given in two panels (with Ai and Aii continuing one after the other) so that protein acronyms and fold changes would be legible. Expression changes of the same proteins in the ADE4/CTRLE4 group are shown alongside for comparison. **B**. Volcano plot of DEPs in the ADE3/CTRLE3 group, using the abundance ratios from PD2.4, which had at least a 20% fold change, the consistent direction of fold-change across the two software platforms (Scaffold and PD2.4), and were identified in 50% or more of subjects. To avoid crowding, we have highlighted only a few DEPs, with a complete list of DEPs shown in the heatmap and shown with greater detail in Table S3. **C**. Heatmap showing a total of 71 proteins differentially expressed, including 37 upregulated and 34 downregulated, in ADE4 relative to E4 controls based on the ADE4/CTRLE4 data shown in Table S2. Expression change of the same proteins in the ADE3/CTRLE3 group are shown alongside for comparison. **D**. Volcano plot of DEPs in ADE4/CTRLE4 was created using the abundance ratios from PD2.4, which had at least a 20% fold change, the consistent direction of fold-change across the two software platforms (Scaffold and PD2.4), and were identified in 50% or more of subjects. To avoid crowding, we have highlighted only a few DEPs, a complete list of DEPs shown in the heatmap, and shown in greater detail in Table S2. **E**. Heatmap showing a total of 51 proteins differentially expressed, including 25 upregulated and 26 downregulated DEPs in ADE4 relative to E3 controls based on the ADE4/CTRLE3 data shown in Table S5. **F**. Volcano plot of DEPs in ADE4/CTRLE3, using the abundance ratios from PD2.4, which had at least a 20% fold change, the consistent direction of fold-change across the two software platforms (Scaffold and PD2.4), and were identified in 50% or more of subjects. To avoid crowding, we have highlighted only a few DEPs, with the complete list of DEPs shown in the heat map, and shown in greater detail in Table S5 **G**. Heatmap showing a total of 104 proteins differentially expressed, including 48 upregulated and 56 downregulated in control ε4 relative to ε3 controls based on the CTRLE4/CTRLE3 data shown in Table S4. The heatmap is split into two panels (Gi and Gii) so that protein acronyms and fold changes would be legible. **H**. Volcano plot of DEPs in CTRLE4/CTRLE3 using the abundance ratios from PD2.4, which had at least a 20% fold change, the consistent direction of fold-change across the two software platforms (Scaffold and PD2.4), and were identified in 50% or more of subjects. To avoid crowding, we have highlighted only a few DEPs, with a complete list of DEPs shown in the heat map and more detail in Table S4.

**Figure 3:**
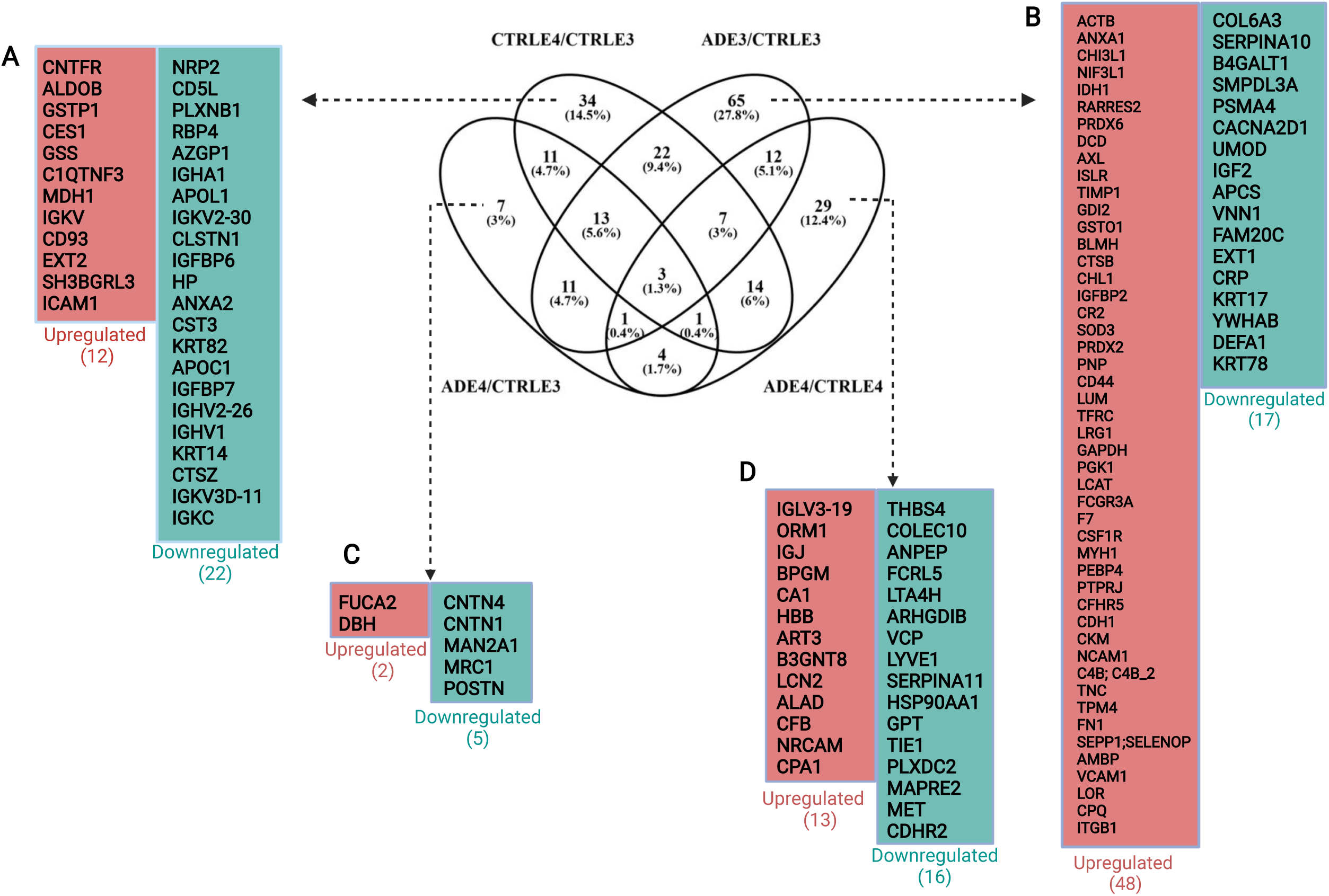
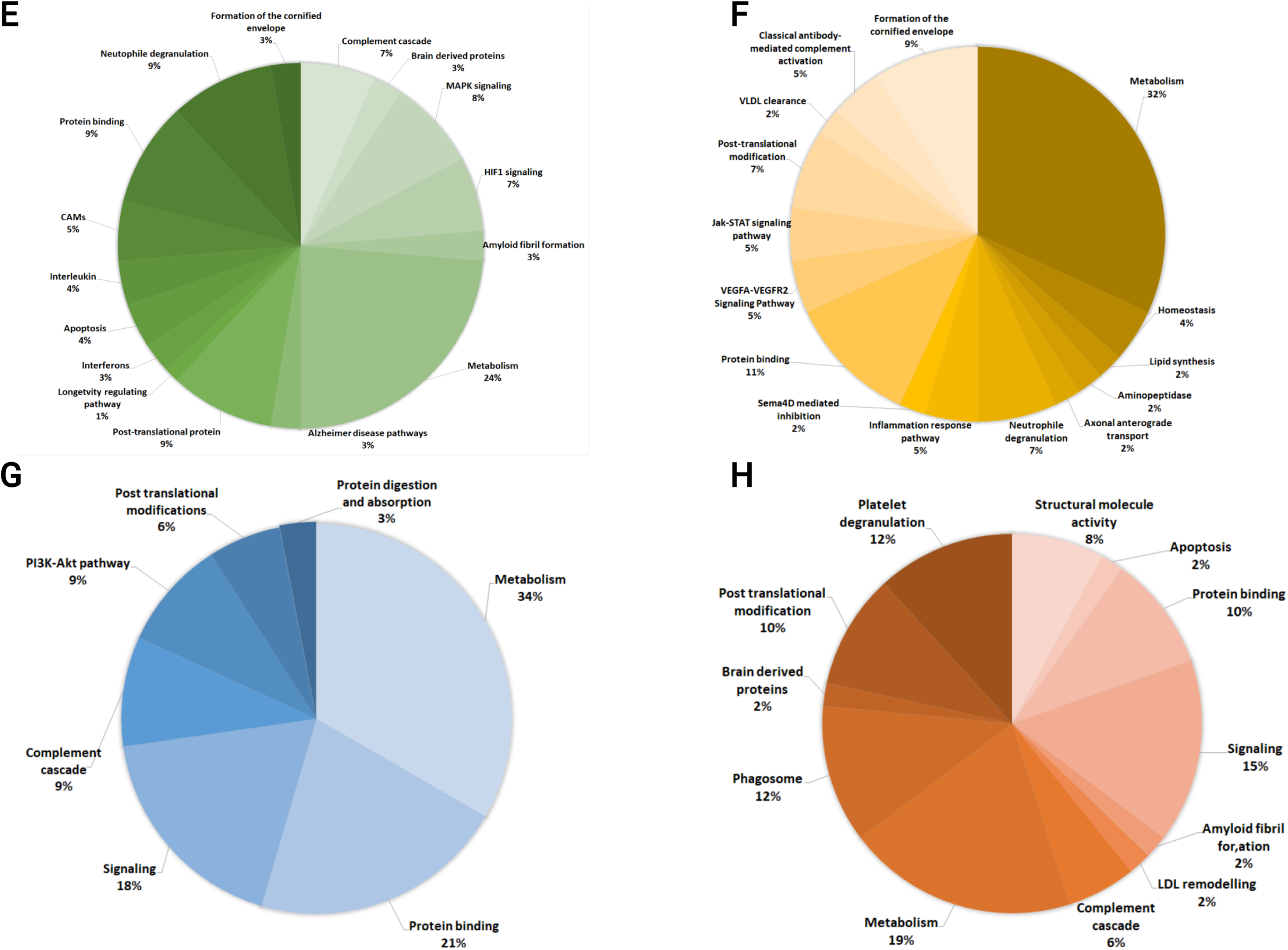
Venn diagram of overlapping and unique DEPs in four groups; CTRLE4/CTRLE3, ADE3/CTRLE3, ADE4/CTRLE3, ADE4/CTRLE4, with DEPs unique to each group displayed on the periphery, as follows; **A**. 34 DEPs unique to the CTRLE4/CTRLE3 group, comprising 12 upregulated and 22 downregulated, in control E4 relative to control E3 (greater detail of the complete list of DEPs identified in the CTRLE4/CTRLE3 group is shown in Table S4). This list contains proteins associated with protection against cognitive decline and neuropathology in APOEε4 carriers who remain cognitively normal. **B**. 65 DEPs (48 upregulated and 17 downregulated) were unique to ADE3/CTRLE3 (complete DEP list and more detail can be found in Table S3). These are DEPs observed in AD subjects who do not carry an APOEε4 allele, so protein expression changes are associated with AD but unrelated to the E4 allele. **C**. 7 DEPs (2 upregulated and 5 downregulated), unique to the ADE4/CTRLE3 group (complete DEP list and more detail can be found in Table S5). These DEPs may reflect the contribution of the E4 allele to AD since the ADE3/CTRLE3 group does not share them. **D**. 29 DEPs (13 were upregulated and 16 downregulated) that were explicitly dysregulated in the ADE4/CTRLE4 group this list represents an experimental correction for the presence of the E4 allele by using normal controls which are heterozygous carriers of the E4 allele (complete DEP list and more detail can be found in Table S2). Pie charts categorizing all the unique DEPs in each group into their biological processes and molecular pathways based on gene ontology (GO) **E**. ADE3/CTRLE3, **F**. CTRLE4/CTRLE3, **G**. ADE4/CTRLE4, **H**. ADE4/CTRLE3.

These unique DEPs were further manually categorized, based on gene ontology, to various biological activities, using data from the PD2.4 analyses (Figure 3E-H). More detailed GO enrichment analysis using STRING software was also performed (Table S7 and Figure S3). The three groups with the most significant proportion of DEPs included metabolism (38%), protein binding (11%), and formation of the cornified envelope (9%) (Figure 3F). Interestingly, from this list, two DEPs, serum amyloid P-component (APCs) and lactotransferrin (LTF), have been linked to the formation of amyloid fibrils^19^. Two other DEPs were linked to Alzheimer’s disease pathway^20,21^: peroxiredoxin-2 (PRDX2) and extracellular superoxide dismutase (SOD3). Other pathways implicated in neurodegenerative disease included; MAPK activation (6 DEPs; Actin cytoplasmic 1 (ACTB), Annexin A1 (ANXA1), voltage-dependent calcium channel subunit alpha-2/delta-1 (CACNA2D1), and 14-3-3 protein beta/alpha (YWHAB), Fibronectin (FN1), and Proteasome subunit alpha type-4 (PSMA4) and HIF1 signalling pathways (5 DEPs; glyceraldehyde-3-phosphate dehydrogenase (GAPDH), insulin-like growth factor-binding protein 2 (IGFBP2), phosphoglycerate kinase 1 (PGK1), transferrin receptor protein 1 (TFRC), and metalloproteinase inhibitor 1 (TIMP1), (Figure 3F and Table S3). Previous studies have demonstrated the role of HIF1 signalling in neurodegenerative disease^22,23^. Given that *APOEε3* is the most common population variant, these DEPs may provide insights into the underlying processes related to AD, but not due to the presence of the *APOEε4* variant.

### Proteome changes seen in *APOEε4* carriers

The 71 DEPs observed in *APOEε4* carriers (ADE4/CTRLE4) included 37 upregulated and 34 downregulated DEPs in ADE4 relative to E4 controls (Table 2, Figure 1E and 2D). The DEPs in ADE4/CTRLE4 are shown in volcano plot format (Figure 2D). A complete list of DEPs in *APOEε4* carriers (ADE4/CTRLE4) is presented in Table S2 and Table 2. Further, GO analysis using STRING was performed to obtain the detailed GO enrichment shown in Table S8 and Figure S3.

When comparing AD *ε4* to control *ε3*, a total of 51 DEPs were identified, including 25 upregulated and 26 downregulated proteins (Figure 1E, Table 2), which are also shown in the heatmap (Figure 2E) and a volcano plot (Figure 2F). A complete list of DEPs for the ADE4/CTRLE4 comparison is presented in Table S5A and Table 2. GO analysis revealed that metabolism (19%), signalling (15%), phagosome (12%), platelet degranulation (12%), and platelet degranulation (10%) were the functional groups containing the majority of ADE4/CTRLE4 DEPs (Figure 3H). Other GO enrichments include amyloid fibril formation (LTF), apoptosis (DSG1), LDL remodelling (CETP), brain-derived neurotrophic factor signalling pathway (PTPRF) (Figure 3H).

While there were some shared DEPs with ADE3/CTRLE3 and ADE4/CTRLE4, we found 29 DEPs (13 upregulated and 16 downregulated) that were uniquely dysregulated in ADE4/CTRLE4 (Figure 3D and Table 2). These AD-related DEPs specific to *ε4* carriers were manually categorized based on gene ontology, with the majority involved in metabolism (34%), protein binding (21%), and signalling (18%). Other enriched categories in ADE4/CTRLE4 included complement cascade, PI3K-Akt pathway, post-translational modifications (PTMs), protein digestion, and protease inhibitors (Figure 3G). The PI3K-Akt pathway, which includes heat shock protein HSP 90-alpha (HSP90AA1), collagen alpha-1(VI) chain (COL6A1), and thrombospondin-4 (THBS4), was one of the distinct pathways dysregulated. Other DEPs included rho GDP-dissociation inhibitor 2 (ARHGDIB) involved in rho GTPase signalling, tyrosine-protein kinase receptor Tie-1 (TIE1) in Rac1/Pak1/p38/MMP-2 pathway, neutrophil gelatinase-associated lipocalin (LCN2) in interleukin-4 and 13 signallings, and lymphatic vessel endothelial hyaluronic acid receptor 1 (LYVE1) in hyaluronan uptake and degradation (Figure 3G and Table S2B).

To explore the role of the *APOE* allele in AD, both ADE4 and ADE3 groups were expressed relative to CTRLE3 (Figure 4D). The majority of the DEPs in these two comparisons were unique to each group (18 in ADE4/CTRLE3 and 87 in ADE3/CTRLE3; Figure 4A), while 28 DEPs were shared, of which the majority (26 DEPs) varied in a similar direction of fold change (Figure 4D, Table S5D). The majority of these DEPs were involved in metabolism (19%), signalling (15%), platelet degranulation (12%), phagosome (12%), and PTM (10%) (Figure 3H).

**Figure 4:**
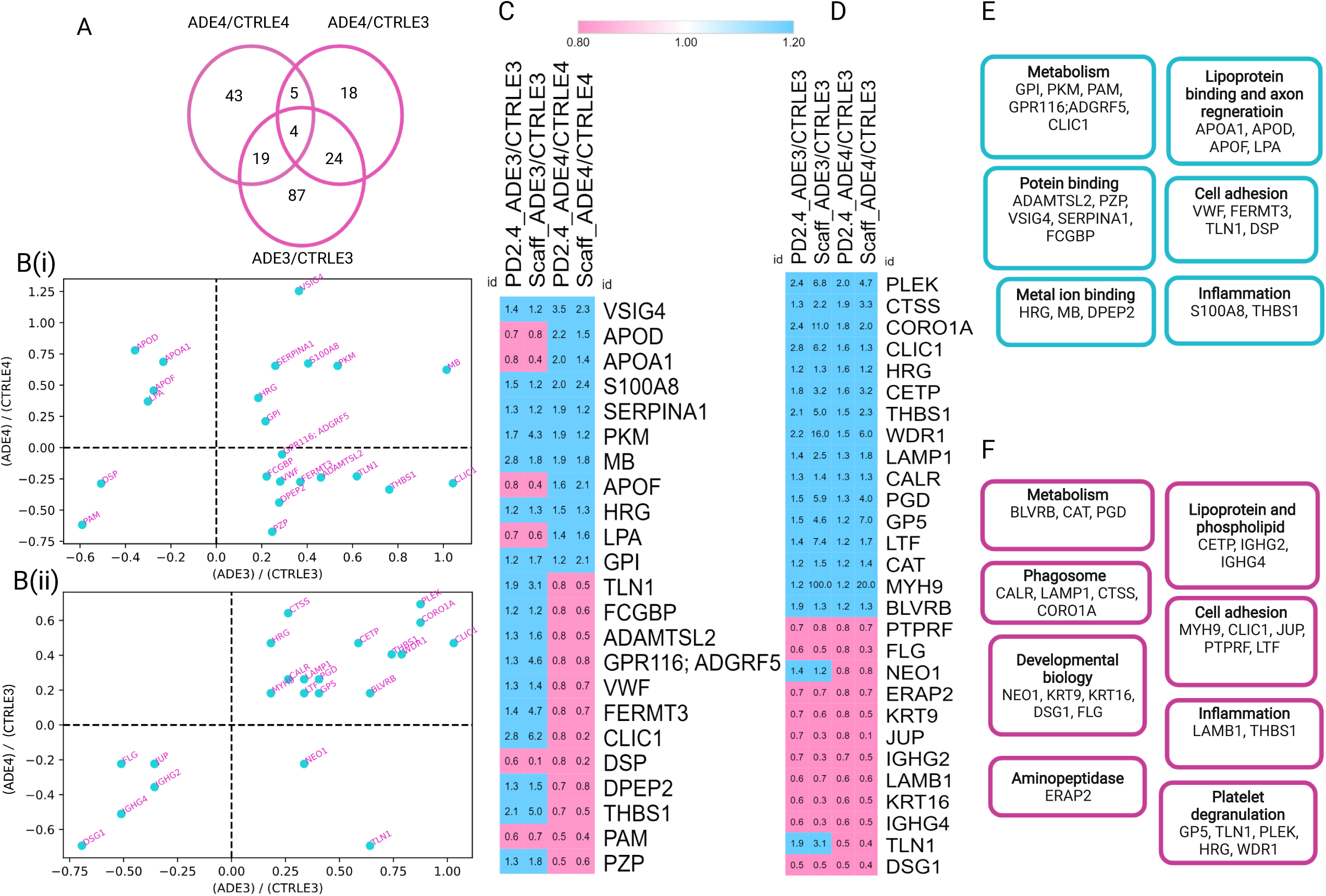
ADE4 and ADE3 expressed relative to E3 homozygous controls and E4 heterozygous carriers to identify proteomic expression changes which are AD specific and those which are contributed by the presence of the E4 allele. **A**. Three-way Venn diagram showing unique and shared DEPs in three AD comparison groups; ADE4/CTRLE4, ADE3/CTRLE3, and ADE4/CTRLE3 **B(i)**. Scatter plot of the DEPs shared in the ADE4/CTRLE4 and ADE3/CTRLE3 groups. **B(ii)**. Scatter plot of the DEPs shared in the ADE4/CTRLE3 and ADE3/CTRLE3 groups. **C**. Heatmap of the 23 DEPs shared in the ADE4/CTRLE4 and ADE3/CTRLE3 groups **D**. Heatmap of the 28 DEPs shared in the ADE4/CTRLE3 and ADE3/CTRLE3 groups. **E**. 23 DEPs common to both ADE4/CTRLE4 and ADE3/CTRLE3 groups are categorized into their biological process based on STRING software. **F**. 28 DEPs common to both ADE4/CTRLE3 and ADE3/CTRLE3 groups, categorized into their biological process based on STRING software.

### Changes common to both *APOE* genotypes in AD (AD risk factors independent of *APOE* allele)

DEPs common to both ADE4/CTRLE4 and ADE3/CTRLE3 groups might indicate AD pathology independent of *APOE* genotypes. The majority of DEPs were specific to each group (43 in ADE4/CTRLE4 and 87 in ADE3/CTRLE3; Figure 4A), while 23 DEPs were common to the ADE4 and ADE3 groups relative to their controls, as indicated in the Venn diagram (Figure 4A), scatter plot (Figure 4B) and heatmap (Figure 4C). Interestingly, only 9/23 DEPs were dysregulated in the same direction in both AD groups (7 elevated in the top right quadrant and 2 downregulated in the bottom left, Figure 4B), whereas 14/23 DEPs were dysregulated in opposite directions. The top left quadrant of Figure 4B shows 4/23 proteins that were elevated in ADE4/CTRLE4 but downregulated in ADE3/CTRLE3. Additionally, 10/23 proteins are shown in the bottom right, indicating downregulated proteins in ADE4/CTRLE4 but were increased in ADE3/CTRLE3. These 23 DEPs were classified into six GO-based functional groups, including metabolism and protein binding, each containing 5 DEPs (Figure 4E). Lipoprotein binding and cell adhesion contained 4 DEPs each. Metal ion binding and inflammation contained 3 and 4 DEPs, respectively. Some GO terms were upregulated in both AD groups, such as innate immune system, glycolysis/gluconeogenesis, myelin sheet, and the complement cascade. On the other hand, some GO terms such as lipid transport, lipid metabolism, lipoprotein metabolic process, cholesterol efflux, and focal adhesion were downregulated in ADE3/CTRLE3 and upregulated in ADE4/CTRLE4. The GO terms contain DEPs of a similar direction of fold change in both groups, suggesting disrupted pathways in AD, irrespective of *APOE* genotype.

Further, we identified 28 DEPs common in ADE4/CTRLE3 and ADE3/CTRLE3 (Figure 4A and Table S5D) were plotted using a scatter plot to show the direction of fold change in common DEPs (Figure 4Bii). 16/28 DEPs were upregulated in both ADE4/CTRLE3 and ADE3/CTRLE3, whereas 10/28 downregulated in both the AD groups. Only 2 DEPs were dysregulated in the opposite direction, i.e., NEO1 and TLN1 were upregulated in ADE3/CTRLE3 whereas downregulated in ADE4/CTRLE3 (Figure 4Bii and 4D, Table S5D). Next, we performed a heatmap using these 28 DEPs showing the PD2.4 abundance ratio and scaffold fold change of each protein in both AD groups (Figure 4D). These common DEPs were further summarized into their functional categories (Figure 4F and Table S5D).

#### Proteins linked to *APOEε4* genotype in controls

A total of 105 proteins were differentially expressed, including 48 upregulated and 56 downregulated proteins (Figure 1E) in control *ε4* relative to control *ε3* (CTRLE4/CTRLE3) are shown in the heatmap Figure 2Gi and 2Gii and Table S4A. In addition, the DEPs in CTRLE4/CTRLE3 are shown in volcano plot format (Figure 2H).

After removing the DEPs common to both AD groups, 34 DEPs unique to CTRLE4/CTRLE3 remained, comprising 12 upregulated and 22 downregulated DEPs (Figure 3A and Table S3B) (Venn diagram). This list of 34 DEPs may provide insight into potential protective mechanisms that prevented these age-matched controls from progressing to AD. These DEPs, unique to control *ε4* carriers, are involved in VLDL clearance (Apolipoprotein C-I, APOC1), lipid synthesis (CD5 antigen-like, CD5L), homeostasis (Complement C1q tumour necrosis factor-related protein 3, C1QTNF3 and SH3 domain-binding glutamic acid-rich-like protein 3 SH3BGRL3), which are essential in maintaining cell integrity. Additional DEPs, unique to control *ε4* carriers, include; Fructose-bisphosphate aldolase B (ALDOB), which is associated with gluconeogenesis; signalling pathways neuropilin-2 (NRP2) and insulin-like growth factor-binding protein 7 (IGFBP7) were downregulated in the VEGFA-VEGFA2 signalling pathway while Annexin A2 (ANXA2) and ciliary neurotrophic factor receptor subunit alpha (CNTFR) are involved in Jak-STAT signalling (Figure 3A). As was the case with ADE4/CTRLE4 and ADE3/CTRLE3, metabolism and protein binding represent biological processes with the most significant proportion of DEPs accounting (Figure 3E).

## Discussion

In this study, both *APOEε3* and *APOEε4* carriers with PiB PET imaging confirmed AD was shown to have a large number of protein expression changes in plasma, with functions including complement cascade, glycolysis, metabolism, plasma lipoprotein assembly, remodelling, and clearance. In addition, several proteins were dysregulated in the presence of the *APOEε4* genotype relative to *APOEε3* in both AD and control groups. This suggests that while some pathways are dysregulated by *APOEε4*, there are shared mechanisms toward developing AD independent of the *APOE* genotype. Furthermore, DEPs unique to *ε4* carriers in the control group suggest potential mechanisms that may protect from progression to AD.

### Plasma level of apolipoproteins and *APOE* genotype

Apolipoproteins are among the most abundant proteins in the brain, with functions relating to cholesterol and lipid transport, and are critical for distributing and recycling lipids in the brain^24^. Differentially expressed apolipoproteins identified in the current study include APOA1, APOA2, APOC1, APOC3, APOD, and APOF, which were downregulated in ADE3/CTRLE3 while APOM, APOA1, APOD, and APOF were upregulated in ADE4/CTRLE4 (Table S2 and S3), and APOB, APOE, APOM, APOD, APOC were upregulated in ADE4/CTRLE3. Conversely, APOD, APOM, and APOA1 were downregulated in the CTRLE4/CTRLE3 group (Table S4i), suggesting a potentially protective effect since these same proteins are upregulated in ADE4. Notably, the level of APOF was unaffected in CTRLE4/CTRLE3, and several other apolipoproteins were downregulated in the CTRLE4/CTRLE3 group, including apolipoproteins LPA, APOL1, APOA2, APOC1, and APOC3. These opposing directions of apolipoproteins expression change in ADE4 and CTRLE4 groups suggest that rather than *APOEε4* producing a similar “toxic effect” in both controls and AD. The response of control vs. AD subjects to the presence of the *APOEε4* allele is qualitatively different and may be the basis of protection from disease progression in CTRLE4, while the ADE4 group succumbs to pathology. A variety of other proteins also have opposite fold change directions in the CTRLE4/CTRLE3 group compared with the ADE4/ADE3 and/or the ADE4/CTRLE4 groups (Table S9). This divergent response of controls and AD subjects to the presence of the *APOEε4* allele explains the paradoxically higher number of DEPs in the *ε4* “corrected” ADE4/CTRLE4 group (71 DEPs) than in the ADE4/CTRLE3 group (51 DEPs), Figures 1 and 4. Surprisingly, by far the longest list of DEPs was in the ADE3/CTRLE3 group (134 DEPs).

That APOC3 was downregulated in ADE3/CTRLE3 and upregulated in ADE4/CTRLE3 suggests that APOC3 may be influenced by the presence of the *APOEε4* allele. Previous work has linked higher APOC3 levels in HDL to an increased risk of coronary heart disease and diabetes, both of which are known risk factors for dementia^25,26^. Higher *APOE* levels in HDL lacking APOC3 in an elderly population were related to better cognitive function and a lower risk of AD dementia^27^. In this context, it is of note that the lipid-binding affinity of *APOEε4* is higher than those of *APOEε2* and *APOEε3*^22,23^, a property that likely accounts for the tendency of *APOEε4* to associate with VLDL, while the *ε2* and *ε3* alleles related to HDL. Such a redistribution of lipoprotein particle composition may also affect expression, half-life, or distribution of other apolipoproteins in *APOEε4* carriers. APOA1 is the principal structural apolipoprotein found in all HDL detectable in the blood. According to Koch et al. 2020, the presence of APOA1 in HDL does not affect the cognitive function or dementia risk, regardless of the presence of APOC3 or APOC3 in HDL^28^. In the current work, APOA1 was lower in ADE3/CTRLE3 but higher in ADE4/CTRLE4, suggestive of an AD-related association. Upregulation of these apolipoproteins in ADE4 may represent a homeostatic response to compensate for the deleterious *APOEε4* allele. The functional groups involved in lipid transport, lipid metabolism, and cholesterol efflux were upregulated in ADE4, whereas all were downregulated in ADE3. Studies have suggested that cholesterol levels in the brain correlate positively with the severity of AD^29^. Elevated lipid metabolism and cholesterol efflux may be a homeostatic response facilitating cholesterol clearance in the ADE4 group^29^.

A gene ontology category enriched in all AD vs. control comparisons was metabolic changes (Figure 3E-H). Both ADE3 and ADE4 showed upregulation of glycolysis/gluconeogenesis-associated proteins such as glycophosphatidylinositol (GPI) and pyruvate kinase muscle (PKM). PKM catalyzes the transfer of phosphoryl groups from phosphoenolpyruvate to ADP generating ATP and pyruvate^30^. Various studies have reported that increased levels of PKM in AD CSF may indicate compensation for mitochondrial dysfunction^31,32^. In this study, GPI and PKM were differentially expressed in both ADE3/CTRLE3 and ADE4/CTRLE4. This was suggestive of an *APOE* allele independent effect, especially as differential expression of these two proteins was not identified in the ADE4/CTRLE3 and CTRLE4/CTRLE3 groups. Several metabolism-related DEPs were unique to ADE3, including upregulation of 6-phosphogluconate dehydrogenase (PGD), peroxiredoxin-6 (PRDX6), isocitrate dehydrogenase, NADP (IDH1) were involved in glutathione metabolism (GSH). Not only is GSH crucial for antioxidant defence in the central nervous system, but it also plays a critical function in preserving the integrity of the blood-brain barrier^33^. As a result, alterations in GSH metabolism may have a greater impact on neurons than on other cell types ^34^. However, clinical research examining the usefulness of boosting antioxidant activity in protecting or restoring cognitive functions in humans, both healthy individuals and clinical AD patients, has generally reported modest efficacy^35^. Even overexpression of the proteins involved in GHS metabolism may be insufficient to prevent/stop the damage caused by AD pathogenesis.

In the ADE4/CTRLE4 group, DEPs were identified, which were previously reported to be differentially expressed in the CSF of AD patients, including bisphosphoglycerate mutase (BPGM) carbonic anhydrase 1 (CA1) activity increased. In contrast, GHS metabolic protein, i.e., aminopeptidase N (ANPEP) activity, decreased in ADE4/CTRLE4^36^. BPGM regulates the 2,3-BPG content in erythrocytes and is a critical regulator of RBC oxygen supply. Increased expression of BPGM in ADE4 implies that RBC energy enzymes are adapted to AD-related changes. Activation of the 2,3-DPG cycle results in an increase in Hb affinity for oxygen, favouring tissue hypoxia^36^.

A total of 9 DEPs were identified in ADE4 compared to both control *ε3* and *ε4* suggestive of AD-related change in *ε4* carriers, maintained even after partial correction using *ε4* controls. This list includes upregulation of glucose-6-phosphate dehydrogenase (G6PD/H6PD) and platelet-activating factor acetyl-hydrolase (PLA2G7). G6PD and complementing antioxidant systems play critical roles in detoxifying reactive oxygen species (ROS). Therefore the concentration of G6PD is crucial in the antioxidant defence mechanism^37^. A recent study by Evlice et al. 2017 reported upregulation of serum G6PD in AD *APOEε3* carriers compared to healthy controls that might protect oxidative stress^38^. The downregulation of G6PD in ADE4 as compared to both control *ε3* and *ε4* carriers in the current data suggests *APOEε4* allele-related compromise of metabolisms/antioxidant defence in AD.

Several markers related to inflammation were identified in both AD groups, including increased S100A8 expression, with the fold change being twice as large in ADE4/CTRLE4 as in ADE3/CTRLE3. Chloride intracellular channel 1 (CLIC1) is another marker of inflammation that was found to be upregulated in ADE4/CTRLE3 and ADE3/CTRLE3 but downregulated in ADE4/CTRLE4, suggestive of an *APOEε4* allele related to change. CLIC1 protein accumulates in peripheral blood mononuclear cells (PBMCs) and is significantly increased in the chronic inflammatory state of the CNS in neurodegenerative disease. Confocal microscopy examination and electrophysiological studies demonstrate the presence of transmembrane CLIC1 in PBMCs from Alzheimer’s disease (AD) patients^39^. This enables the use of blood tests and other conventional technologies to distinguish between healthy persons and those who are undergoing neurodegenerative processes.

We found upregulation of NEO1 and NCAM1 in ADE3 carriers but no change in ADE4 compared to their respective controls. Neuronal damage markers such as Hepatocyte growth factor receptor (MET) decreased in ADE4/CTRLE4 but was not differentially expressed in ADE3/CTRLE3. The protein CHI3L1 (also called YKL40) is a well-studied CSF protein associated with reactive astrocytes, and in the current work was higher in ADE3/CTRLE3 but unchanged in ADE4/CTRLE3 and ADE4/CTRLE4^40^.

### AD plasma proteomics in *APOEε3* and *APOEε4* carriers

The *APOEε4* allele is the most explored and familiar genetic risk factor for late-onset AD^1^, increasing the risk of AD, as well as the severity and heterogeneity of the pathology^41-43^. However, it is neither an essential nor a sufficient factor for progression to AD since non-carriers of the *ε4* allele also succumb to AD, while many *ε4* carriers do not progress to AD. Comparing AD *ε3* and *ε4* carriers with their respective *ε3* and *ε4* controls may provide insight into *APOE* allele independent proteomic associations with AD, while the same comparison using *ε3* controls only may provide insight into the specific contribution of the *APOEε4* allele to the AD plasma proteome. Though it should be noted that experimental correction with normal controls who are carriers of the *ε3* and *ε4* alleles may not be perfect, since *(1)* the effects of *APOE* alleles may play out differently in AD vs. normal controls, and *(2)* the *ε4* controls, in this case, were all heterozygous, while the AD *ε4* carriers were all homozygous.

The proteins PRDX2 and SOD3 are antioxidant proteins directly linked to Alzheimer’s disease pathway^20,21^ and were uniquely upregulated in AD *ε3* compared to control *ε3*. PRDX2, prevalent in erythrocytes, has been demonstrated to play a critical function in protecting erythrocytes from oxidative stress by scavenging ROS and contributing to cell signalling^44^. Studies have suggested that PRDX2 exists in a more oxidized state in the AD brain than controls^45^. Prx expression is increased, and the ability to retain Prxs at a decreased level is part of a unique neuroprotective process that occurs in response to Aβ build-up^45^. Favirn et al., 2013, investigated some genes that were consistently overexpressed in Aβ *Drosophila* (fruitflies) AD models and identified SOD3 as an Aβ toxicity modifier. They suggested that imbalance of this enzyme may result in an elevated level of the strong oxidant H2O2 in Aβ flies, hence contributing to AD pathology^46^. The PI3K-Akt signalling pathway component collagen alpha-1(VI) (COL6A1) was decreased in ADE4 when compared to *ε3* and *ε4* controls, whereas THBS1 was decreased in ADE4/CTRLE3 and increased in ADE4 when compared *ε4* controls. Reducing collagen VI increased Aβ neurotoxicity, but treating neurons with soluble collagen VI inhibited the attachment of Aβ oligomers with neurons, increased Aβ aggregation, and avoided neurotoxicity^47^. Collagen VI is identified as a critical component of the neural damage response, and its neuroprotective potential has been demonstrated^47^. The downregulation of these proteins uniquely in ADE4 individuals might explain the severity of the disease in *APOEε4* carriers.

The complement system is a major part of the innate immune system, and its classical activation pathway can be directly triggered by amyloid aggregates^48,49^. The involvement of different complement proteins in different cognitive stages suggests that triggers of the complement system may exist that are dependent on the degree of neuronal injury and/or amyloid fibril production. Previous studies have demonstrated upregulation of components of the complement system in the AD brain and the influence of the complement cascade in synapse dysfunction and loss in a mouse model of tauopathy^50,51^. Upregulation of CFB, IGLV3-19, and downregulation of COLEC10 was uniquely identified in ADE4/CTRLE4. Comparing ADE4 with control *ε3* and *ε4*, endothelial protein C receptor (PROCR) was found to be upregulated in both comparisons. Previous studies investigating complement-related protein concentrations in CSF reported divergent results with higher concentrations in AD-type dementia patients^52-54^. Notably, neuroinflammation is more severe in *APOEε4* carriers and related animal model studies^55^, including co-localization of *APOE* with microglia in the brain, implying that *APOE* plays a role in the innate immune response in AD brain^3^. Future research should focus on longitudinal changes in complement levels that occur during the development of AD, as well as the effect of the *APOE* genotype on these processes.

### Differential protein expression in normal controls carrying *APOEε3* and *ε4* alleles

While the *APOEε4* allele is a well-known risk factor for AD, not all who carry this allele progress to AD. Comparing the plasma proteomes of *ε4* and *ε3*carriers in normal controls may provide some insight into factors that provide protection from progression to AD despite the presence of the *ε4* allele. There were 14 DEPs in CTRLE4/CTRLE3 involved in metabolism, showing that dysregulation of metabolism may be a general mechanism of aging rather than a feature of AD (Table S4ii). Glycolysis is required for a range of brain functions, including energy production, synaptic transmission, and redox balance. In both preclinical and clinical AD patients, decreased glycolytic flux has been demonstrated to correlate with the severity of amyloid and tau pathology^56^. Upregulation of glycolysis/gluconeogenesis-related proteins, i.e., ALDOB and GOT1 in control *ε4* compared to control *ε3*, might suggest the protective mechanism increasing the glycolysis metabolism. These metabolic changes may act as a risk indication rather than an independent risk factor. However, specific metabolism markers such as GPI and PKM may help distinguish AD from age-matched controls. A better knowledge of the link between AD and metabolism, as well as how this relationship is modulated by *APOEε*4, will also be necessary.

On the other hand, ALDOB and GOT1 might provide insights into age-matched controls’ protective mechanisms. VLDL clearance, VEGFA signalling, and JAK-STAT pathways were all uniquely enriched in the case of CTRLE4/CTRLE3. Both NRP2 and IGFBP7 were downregulated in the CTRLE4/CTRLE3 group, and both are involved in VEGFA signalling. Despite the complexity and mixed evidence of VEGF associations with AD, there is growing evidence that VEGF may have a neuroprotective role^57^. The VLDL clearance pathway involving APOC1 was differentially expressed in CTRLE4/CTRLE3. APOC1 is predominantly expressed in the liver and is activated during the differentiation of monocytes into macrophages required for HDL and VLDL metabolism. APOC1 has been implicated in various malignancies, and other research points to a link between APOC1 and human longevity^58,59^. Given the discrepancy of research findings, it is critical to discover the role of these pathways in human longevity and healthy aging.

## Limitations

Several limitations of this study include the small sample size, individuals with AD were all homozygous *APOE* ε*4/4* carriers, whereas age-matched controls were withal heterozygous *APOE* ε*3/4* carriers and the absence of confirmation using another technique such as ELISA. In addition, as this is an exploratory study, additional research into the relevance of these proteins is warranted in prospective studies of dementia-free individuals in midlife and long-term dementia incidence follow-up.

## Conclusion

This study performed an in-depth proteome analysis to identify plasma proteome signatures associated with *APOEε3* and *APOEε4*. In late-onset AD, the *APOEε4* allele is the most well-known genetic risk factor. However, non-carriers of the *ε4* allele also succumb to AD, but many *ε4* carriers do not. We identified a high number of protein expression alterations in plasma which were found uniquely in *APOEε3* and *APOEε4* carriers. Interestingly, several proteins were also dysregulated in the presence of both *APOEε3* and *APOEε4* genotypes depicting the involvement of these proteins in the pathogenesis of AD regardless of the *APOE* genotypes. Furthermore, our findings also identified some proteins previously discovered in AD CSF and brain proteomics signatures that could provide clinically meaningful information.

## Supporting information

The overall similarity of low abundance proteins across samples is also evident by SDS PAGE (Supplementary Figure S1)

We identified 1,055 proteins (false discovery rate <1%) with 23,242 total peptides using Proteome Discoverer 2.4 (PD2.4) search engine and 800 protein

## Captions

**Table S1:** Total number of proteins identified in PD2.4 and scaffold search engines in all 3 comparisons. Table S1A; Total number of confidently identified plasma proteins in ADE3/CTRLE3, ADE4/CTRLE4, and CTRLE4/CTRLE3 using Proteome Discoverer 2.4. Table S1B; Total number of confidently identified plasma proteins in ADE3/CTRLE3Scaffold Q+ software v 4.11.0. Table S1C; Total number of confidently identified plasma proteins in ADE4/CTRLE4Scaffold Q+ software v 4.11.0.

**Table S2:** List of differentially expressed proteins (DEPs) in the ADE4/CTRLE4 group, based on the following selection criteria; quantified in >5 individuals, proteins identified with a minimum of two peptides, the consistent direction of protein fold change across two bioinformatics platforms with orthogonal quantification approaches (peak area ratio with PD2.4 and spectral counting with Scaffold) with a fold change of at least 20% (≤0.08 and ≥ 1.2) in both search engines. **Table S2A**: 71 total DEPs (37 upregulated and 34 downregulated), identified in the AD E4/CTRLE4 group. **Table S2B:** 29 DEPs unique to theAD4/CTRLE4 group, i.e., not meeting selection criteria in the ADE4/CTRLE3, ADE3/CTRLE3, and CTRLE4/CTRLE3 groups, so are considered the DEPs unique to the ADE4/CTRLE4 group.

**Table S3:** List of differentially expressed proteins (DEPs) in the ADE3/CTRLE3 group, based on the following selection criteria; quantified in >5 individuals, proteins identified with a minimum of two peptides, the consistent direction of protein fold change across two bioinformatics platforms with orthogonal quantification approaches (peak area ratio with PD2.4 and spectral counting with Scaffold) with a fold change of at least 20% (≤0.08 and ≥ 1.2) in both search engines. **Table S3A:** 134 total DEPs (48 upregulated and 56 downregulated) were identified in the ADE3/CTRLE3 group. **Table S3B:** 65 DEPs unique to the ADE3/CTRLE3 group. These DEPs did not meet the selection criteria in the ADE4/CTRLE4, CTRLE4/CTRLE3, and ADE4/CTRLE3 groups, so they are considered the DEPs unique to the ADE3/CTRLE3 group.

**Table S4A:** 105 total DEPs (48 upregulated and 56 downregulated), identified in the CTRLE4/CTRLE3 group. **Table S4B:** 34 unique DEPs identified in the CTRLE4/CTRLE3 group. These DEPs did not meet the selection criteria in the ADE3/CTRLE3, ADE4/CTRLE3, and ADE4/CTRLE4 groups, so they are considered the DEPs unique to the CTRLE4/CTRLE3 group.

**Table S5:** List of differentially expressed proteins (DEPs) in the ADE4/CTRLE3 group which meet the following selection criteria; quantified in >5 individuals, proteins identified with a minimum of two peptides, the consistent direction of protein fold change across two bioinformatics platforms with orthogonal quantification approaches (peak area ratio with PD2.4 and spectral counting with Scaffold) with a fold change of at least 20% (≤0.08 and ≥ 1.2) in both search engines. **Table S5A:** 51 DEPs identified in the ADE4/CTRLE3 group. **Table S5B:** 18 DEPs unique to the ADE4/CTRLE3 group. These DEPs did not meet the above selection criteria in the ADE3/CTRLE3 or ADE4/CTRLE4 groups. **Table S5C:** 9 DEPs identified in the ADE4/CTRLE4 and ADE4/CTRLE3 groups. **Table S5D:** 28 DEPs were identified in the ADE3/CTRLE3 and ADE4/CTRLE3 groups.

**Table S6:** List of differentially expressed proteins (DEPs) identified in the ADE4/ADE3 group, which meet the following criteria; quantified in >5 individuals, proteins identified with a minimum of two peptides, the consistent direction of protein fold change across two bioinformatics platforms with orthogonal quantification approaches (peak area ratio with PD2.4 and spectral counting with Scaffold) with a fold change of at least 20% (≤0.08 and ≥ 1.2) in both search engines. In total, 93 DEPs were identified in the ADE4/ADE3 group.

**Table S7:** GO term enrichment was performed in STRING software using 134 DEPs identified in the ADE3/CTRLE3 group, 95 of which were upregulated and 39 downregulated. This analysis provides insight into the biological processes, cellular components, molecular functions, KEGG & reactome molecular pathways affected in the ADE3/CTRLE3 group.

**Table S8:** GO term enrichment was performed in STRING software using 71 DEPs identified in the ADE4/CTRLE4 group, 37 of which were upregulated and 34 of which were downregulated. This analysis provides insight into the biological processes, cellular components, molecular functions, KEGG & reactome molecular pathways affected in the ADE4/CTRLE4 group.

**Table S9:** This table contains the list of differentially expressed proteins (DEPs) those quantified in >5 individuals, proteins identified with a minimum of two peptides, the consistent direction of protein fold change across two bioinformatics platforms with orthogonal quantification approaches (peak area ratio with PD2.4 and spectral counting with Scaffold) with a fold change of at least 20% (≤0.08 and ≥ 1.2) in both search engines.

A list of all DEPs identified opposite fold change directions in the CTRLE4/CTRLE3 group compared with the ADE4/ADE3 and/or the ADE4/CTRLE4 groups.

