## Supplementary material for "Impact of APOE ε3 and ε4 genotypes on plasma proteome signatures in Alzheimer’s disease": The overall similarity of low abundance proteins across samples is also evident by SDS PAGE (Supplementary Figure S1)

**Figure S1:** Images of NuPAGE LDS gels of depleted plasma samples from individual subjects containing low abundant plasma proteins (LAP), which was the flow-through from the HU14 column. Each gel lane contained an equal loading of total protein (50 ug total proteins were loaded per gel lane). We have used 10 LAP samples from each subject; A3 denotes ADE3, A4 denotes ADE4, C3 denotes control E3, and C4 denotes control E4.

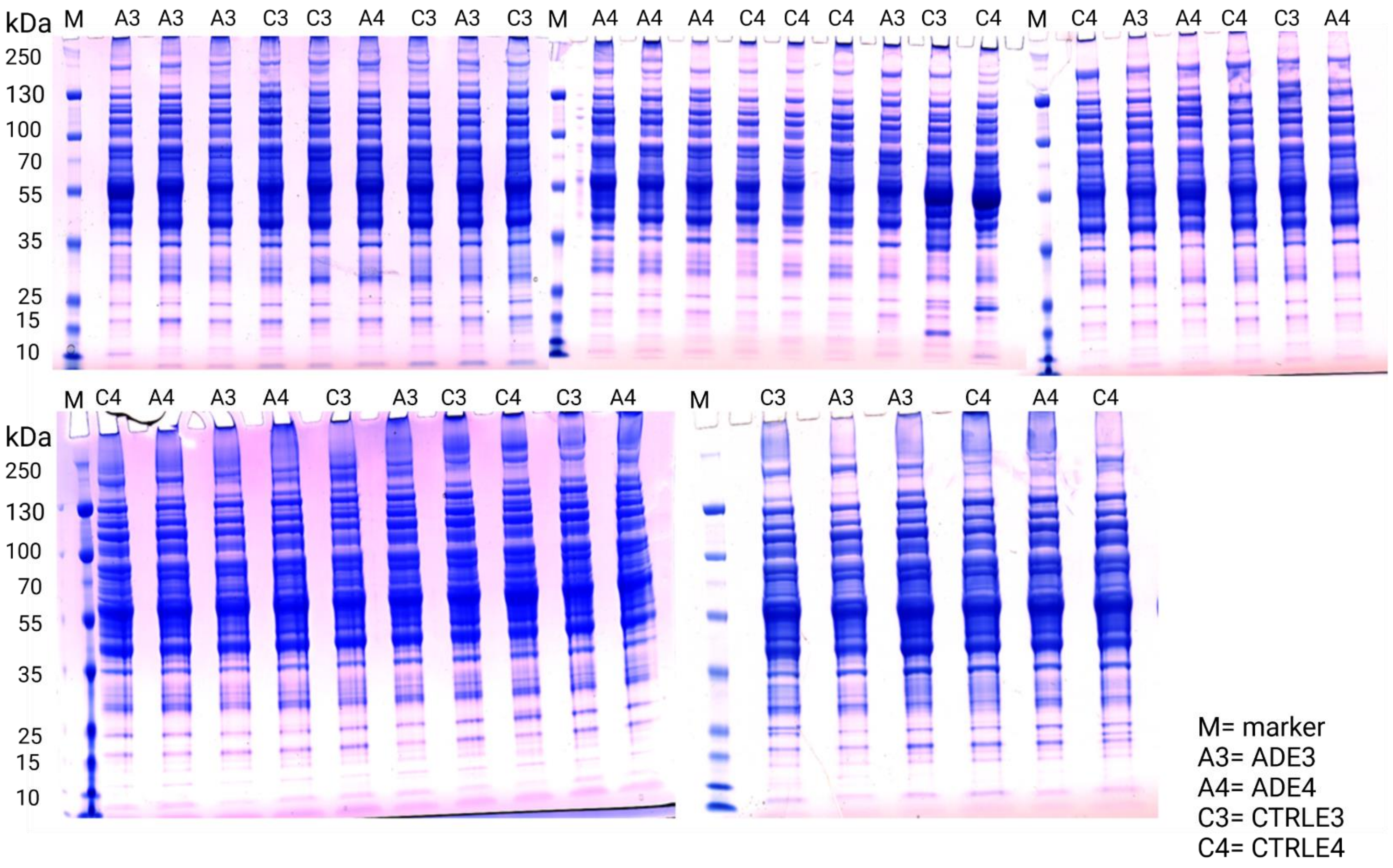

**Figure S2:** Scatter plots comparing PD2.4 versus scaffold fold-change are displayed. Density plots to compare the data points between *APOE* genotypes and respective controls. Each dot represents the abundance ratio of each protein, and the color shows the dot density (**panels A – D**). Scatter plots were also plotted using only DEPs from each comparison. **E.** 134 DEPs from ADE3/CTRLE3 (Table S3), **F.** 51 DEPs from ADE4/CTRLE3 (Table S5), **G.** 93 DEPs from ADE4/ADE3 (Table S6), **H.** 105 DEPs from CTRLE4/CTRLE3 (Table S4), **I.** 71 DEPs from ADE4/CTRLE4 (Table S2).

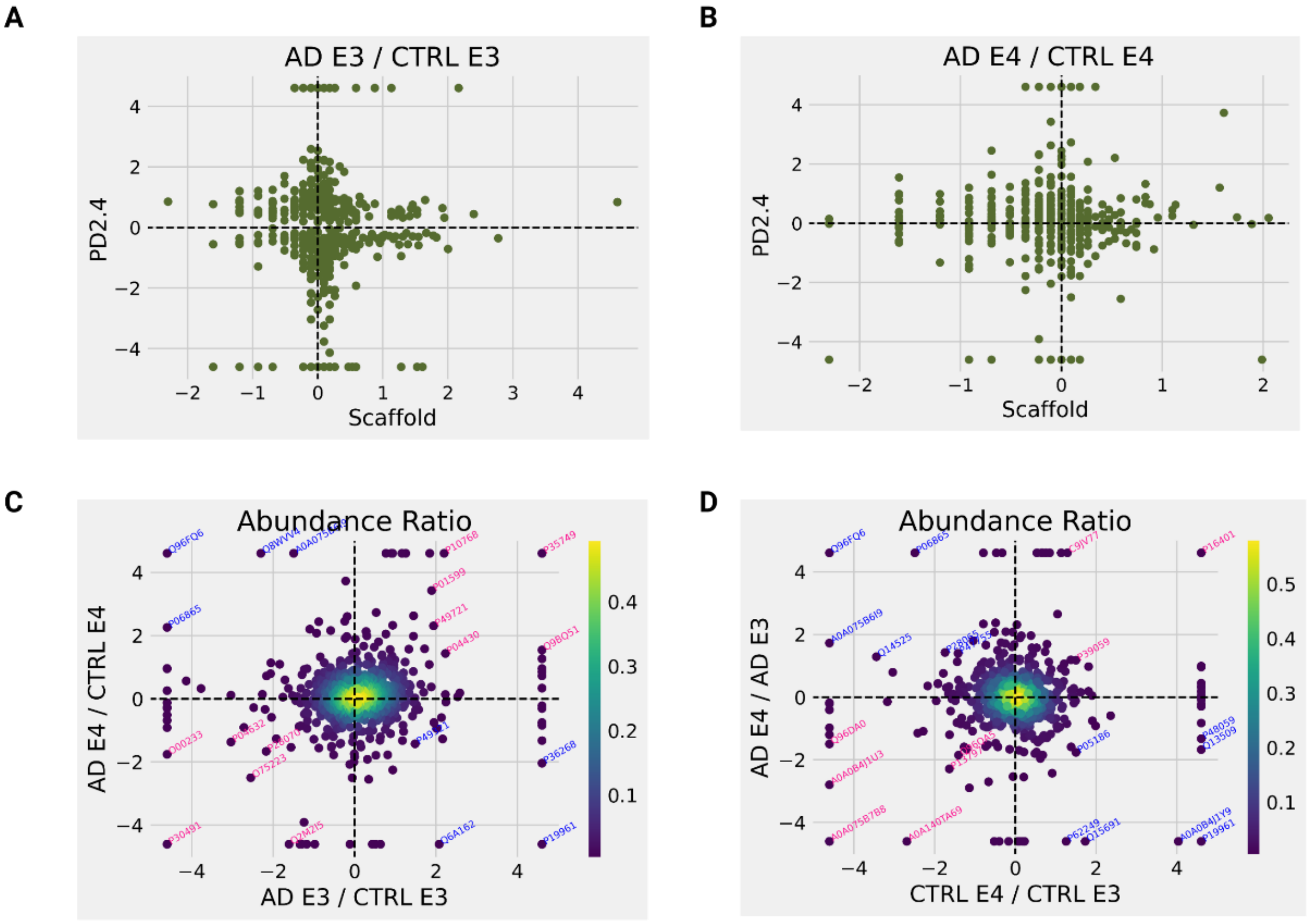

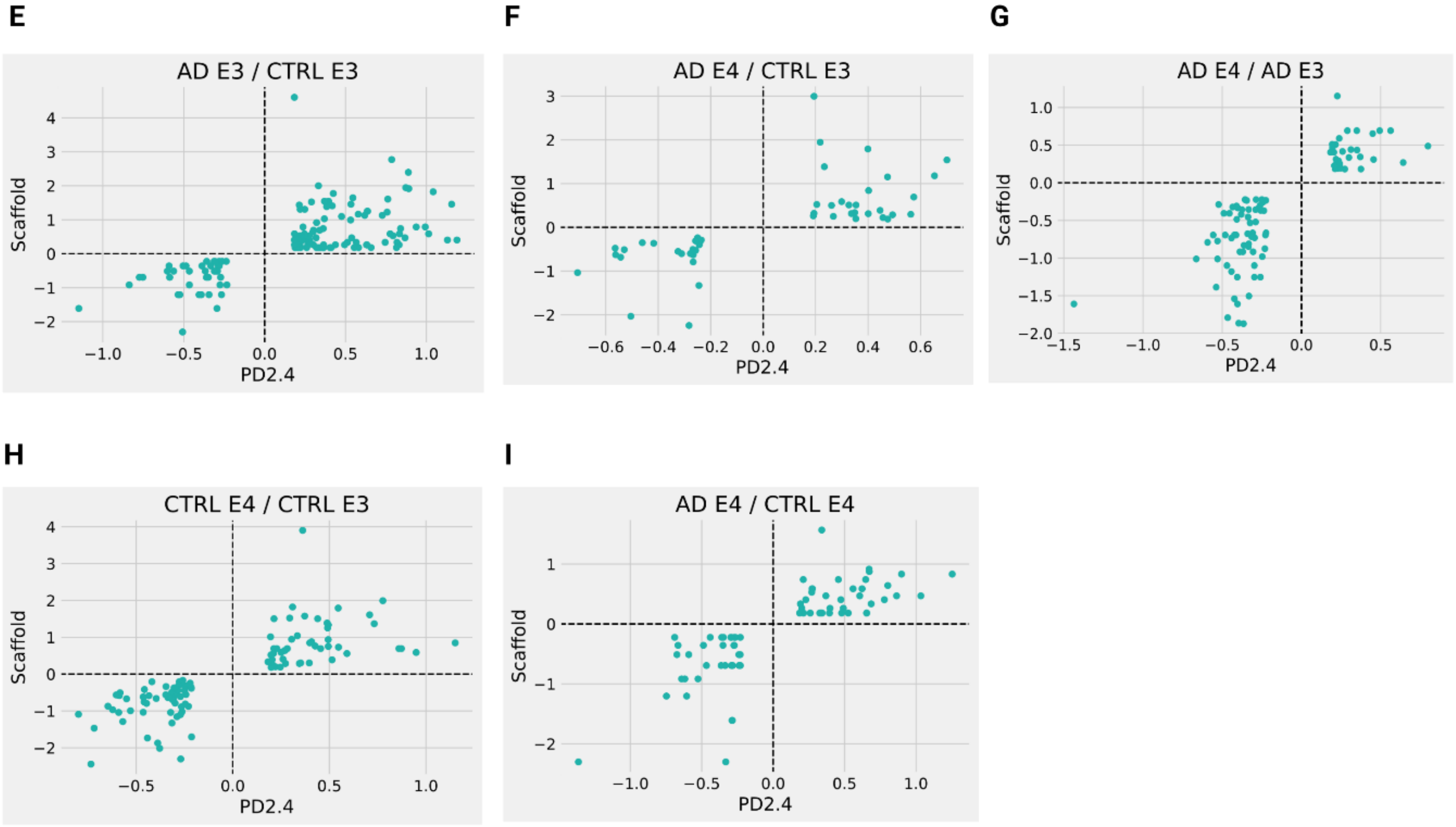
E
AD E3 / CTRL E3
Scaffold
PD2.4
F
AD E4 / CTRL E3
Scaffold
PD2.4
G
AD E4 / AD E3
Scaffold
PD2.4
H
CTRL E4 / CTRL E3
Scaffold
PD2.4
I
AD E4 / CTRL E4
Scaffold
PD2.4

**Figure S3:** Gene ontology enrichments common to the ADE3/CTRLE3 and ADE4/CTRLE4 groups to understand the significantly dysregulated pathways. DAVID v6.8 software was used for enrichment analysis, and the data used in the graphical displays were prepared manually sorting from PD2.4 analyses. **A.** biological processes, **B.** molecular function, **C.** cellular components, and **D.** KEGG pathways. This figure presents a complete list of GO term enrichments, using DEPs commonly identified in both AD groups (details of the DEPs used are shown in Table S7 and S8). These pathways were identified using the list of DEPs common to both ADE4 and ADE3 groups, suggesting that these pathways are disrupted in AD irrespective of the genotype. **SampleGroup:** Red circle- GO upregulated in ADE4; Blue plus- GO downregulated in ADE4; Green square- GO upregulated in ADE3; Yellow triangle- GO downregulated in ADE3. **Count:** The size of the symbols represents the number of DEPs involved in each GO enrichment term. Apart from common GO in both AD groups, the complete list of GO in Table S5 contains ADE3 and ADE4 in Table S6.

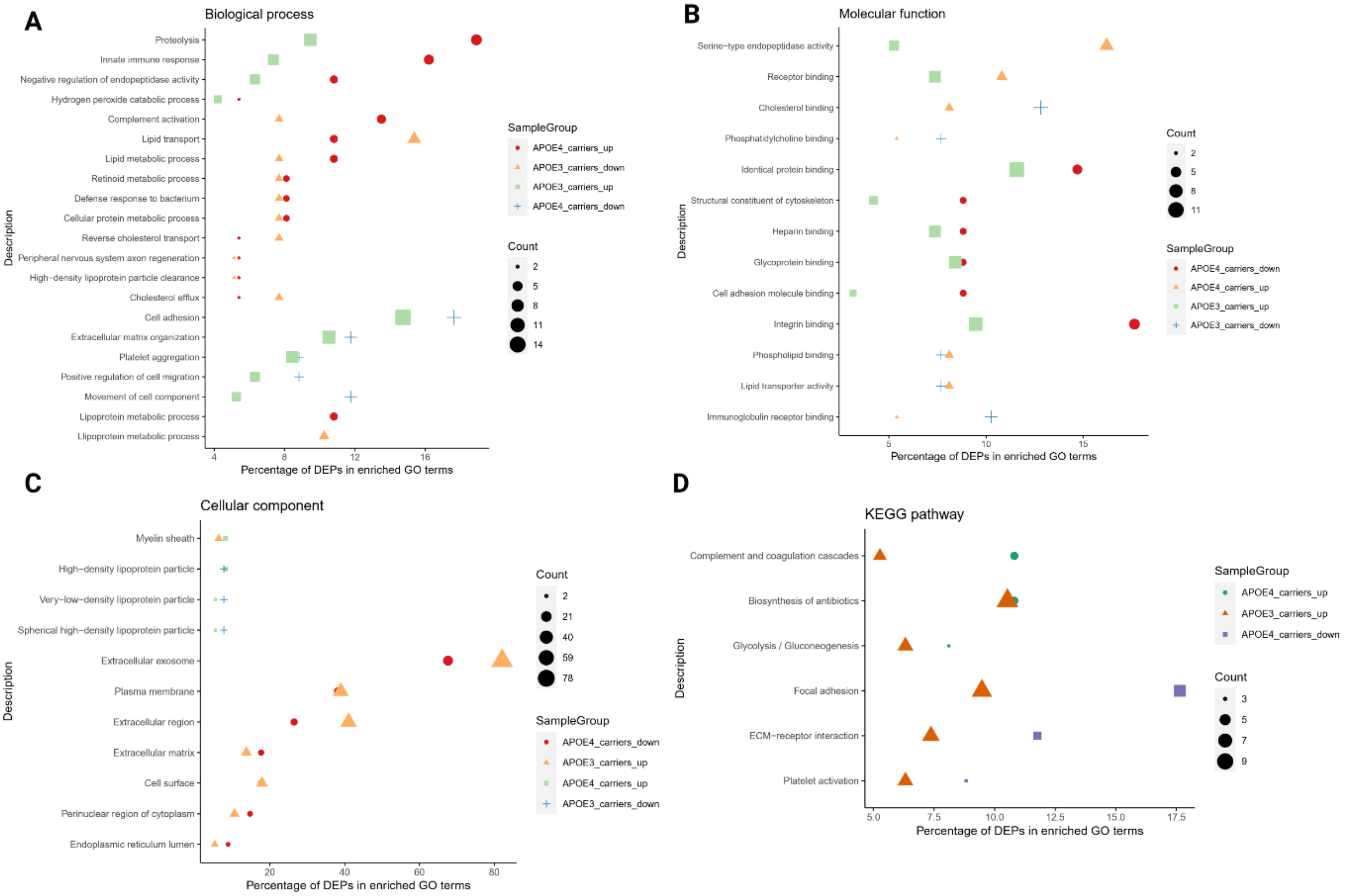
